# A Scribble/Cdep/Rac pathway regulates follower cell crawling and cluster cohesion during collective border cell migration

**DOI:** 10.1101/2022.01.04.474957

**Authors:** Joseph P. Campanale, James A. Mondo, Denise J. Montell

## Abstract

Collective cell movements drive normal development and metastasis. *Drosophila* border cells move as a cluster of 6-10 cells, where the role of the Rac GTPase in migration was first established. Rac stimulates leading edge protrusions in most migratory cells. Upstream Rac regulators in leading border cell protrusions have been identified; however the regulation and function of Rac in follower cells is unknown. Here we show that Rac is required in all cells of the cluster and promotes follower cell motility. We identify a Rac guanine nucleotide exchange factor, Cdep, that also regulates follower cell movement and cluster cohesion. The tumor suppressors Scribble, Discs Large, and Lethal Giant Larva localize Cdep basolaterally and share phenotypes with Cdep. Relocalization of Cdep::GFP partially rescues Scrib knockdown, suggesting that Cdep is a major downstream effector of basolateral proteins. Thus, a Scrib/Cdep/Rac pathway promotes cell crawling and coordinated, collective migration *in vivo*.

## Introduction

Collective cell migrations drive homeostasis, normal development, and tumor metastasis (Friedl and Gilmour, 2009; Friedl et al., 2012; Mishra et al., 2019a; Scarpa and Mayor, 2016; Stuelten et al., 2017). Within collectives, leaders and followers can be distinguished by their positions, and cells at the front steer the group (Khalil and de Rooij, 2019; Revenu et al., 2014; Zhang et al., 2019). The border cell cluster in the *Drosophila* ovary has emerged as a powerful model for elucidating the cellular and molecular mechanisms regulating collective migration (Montell et al., 2012; Norden and Lecaudey, 2019; SenGupta et al., 2021). *Drosophila* egg chambers consist of 15 nurse cells and one oocyte surrounded by a monolayer of follicle cells (Fig. 1A). Follicle cells exhibit classical epithelial polarity with apical microvilli, E-cadherin rich adherens junctions, lateral membranes, and basal surfaces, which contact a basement membrane that surrounds each follicle (Fig. 1A). During developmental stage 9, 6-10 anterior follicle cells called border cells delaminate as a cluster, detach from the basement membrane and their epithelial neighbors, retain apicobasal polarity, and squeeze between the nurse cells to migrate to the oocyte (Montell et al., 2012)[reviewed in] (Fig. 1A).

**Figure 1:**
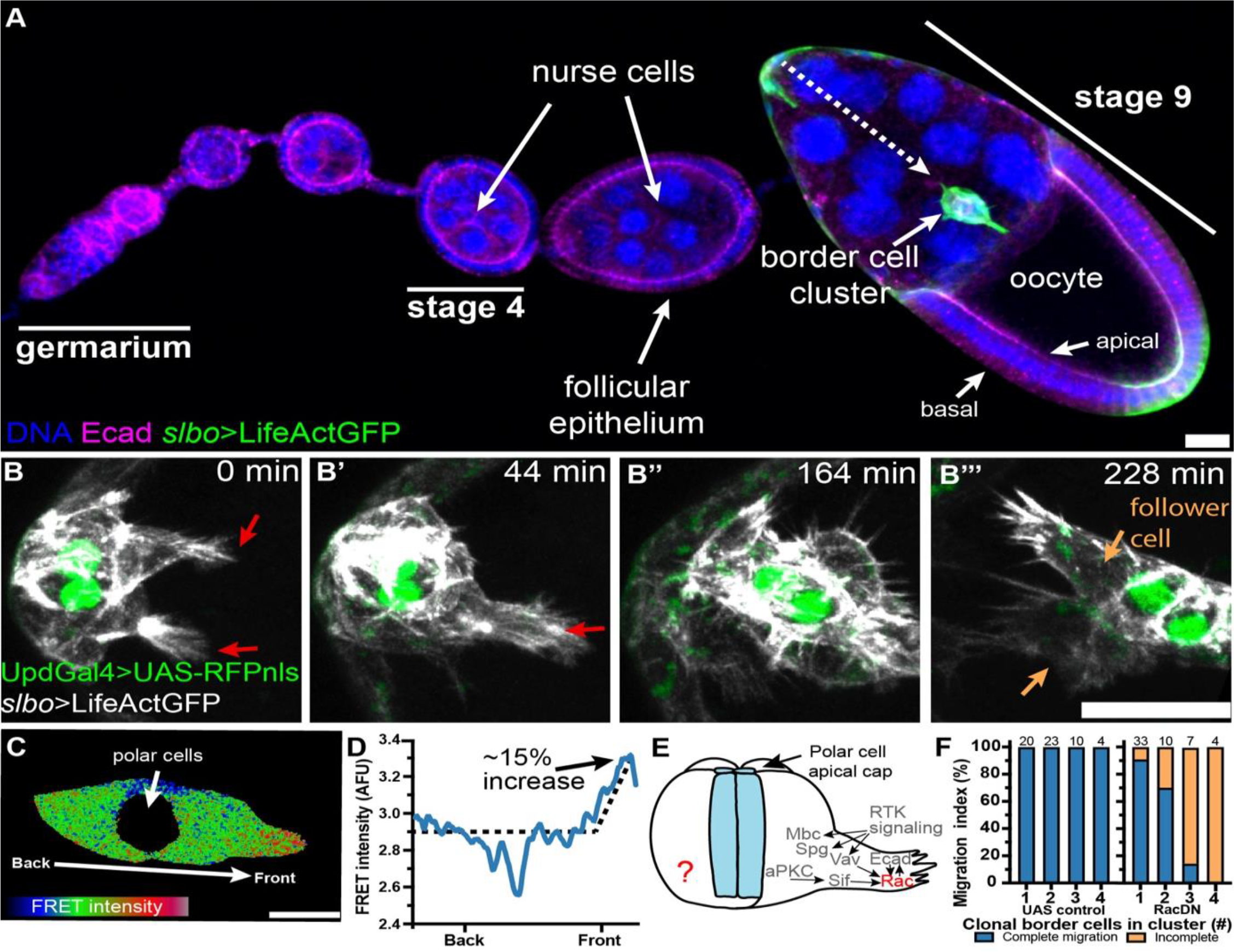
Rac is required in follower cells. (A) Projected *Drosophila* ovariole from ovary with DNA (blue), Ecad (magenta), and *slbo*>LifeActinGFP (green). (B-B’’’) Stills from a time-lapse of delaminating border cells expressing *slbo*>LifeActGFP (white) and polar cell nuclei expressing *upd*Gal4>UAS-RFPnls (green). Lead cell protrusions and follower cells are marked by red and orange arrows, respectively. (C) Intensity coded RacFRET signal co-expressing control UAS-LacZ. (D) Line plot of RacFRET intensity for cluster in (C). (E) Border cell cluster schematic overlayed with known Rac signaling in lead cell protrusions. (F) Frequency distribution for complete border cell migration of clusters containing variable numbers of clonal UAS expressing cells. All scale bars are 20 µm.

The evolutionarily conserved Rho family of small GTPases, Rac, Cdc42 and Rho, regulate the actin cytoskeleton dynamics that drive single and collective cell migrations *in vivo* and *in vitro* (Lyda et al., 2019; Nobes and Hall, 1995; Parri and Chiarugi, 2010; Ridley, 2015). The *Drosophila* genome encodes three Rac genes, Rac1, Rac2 and Mig-2-like (Mtl), which in border cells promote lead cell protrusion and migration (Murphy and Montell, 1996; Geisbrecht and Montell, 2004). Rac activity is highest in the lead cell, and photoactivation of Rac in one cell is sufficient to make it the leader and steer the whole cluster (Wang et al., 2010).

Decades of research have focused on Rac activity and function in lead cell protrusions (Bianco et al., 2007; Duchek et al., 2001; Fernández-Espartero et al., 2013; Geisbrecht and Montell, 2004; Murphy and Montell, 1996; Ramel et al., 2013). These large protrusions have at least two functions. First, they steer the cluster by sensing chemoattractants as well as small open spaces between nurse cells (Dai et al., 2020) and second, they facilitate mechanical communication by inhibiting large protrusions in follower cells (Cai et al., 2014). However the contribution of follower cells to cluster motility and the regulation and function of Rac in follower cells has not been addressed. Follower cell behavior is generally a less well understood feature of collective cell migration, particularly *in vivo (Qin et al>., 2021)*.

Here we show that Rac is required in all cells of the border cell cluster and identify a basally localized Rac GEF, Cdep, that is required for follower cell crawling. We further show that Cdep membrane localization requires the basolateral proteins Scribble (Scrib), Discs Large (Dlg) and Lethal Giant Larva (LGL). Inhibition of any of these components leads to defective cell motility and reduced cluster cohesion. These phenotypes are complementary to knockdown of E-cadherin in the migratory cells, which causes loss of lead cell protrusion but maintenance of cluster crawling and cohesion. This work suggests an integrated model of collective border cell migration in which Rac-dependent lead cell protrusions steer the cluster whereas Rac-dependent crawling in all cells is required for coordinated and cohesive movement.

## Results

### Rac is active and required in follower cells

To investigate the contribution of follower border cells to cluster migration, we performed timelapse imaging of delaminating clusters using markers for the two cell types that make up the cluster: a central pair of non-migratory polar cells, surrounded and carried by migratory cells [reviewed in (Montell et al., 2012) (Fig. 1B-B’’’, Movie 1). Live imaging of border cell F-actin at high spatial and temporal resolution showed that as the migratory cells round up and encase the polar cells, multiple cells generate large forward directed protrusions (Fig. 1B, Movie 1) and appear to compete to lead the cluster. Eventually, one cell “wins” (Fig. 1B’), and both the leader and followers move away from the epithelium (Fig. 1B’-1B’’’).

The dominant negative form of Rac, Rac^T17N^ (RacDN), inhibits all three, functionally redundant Racs in *Drosophila*. Expressing RacDN in all outer migratory cells blocks lead cell protrusion, cluster delamination and migration so the cells remain at the anterior tip of the egg chamber (Murphy and Montell, 1996; Geisbrecht and Montell, 2004). Using a RacFRET activity reporter, we observed that although Rac activity in the lead cell is on average 15% higher than the rear, Rac is also active in follower cells (Fig. 1C and D). However its role in followers, if any, has not been addressed (Fig. 1E). Similarly, the regulatory network of guanine exchange factors (GEFs) that activate Rac in the lead cell have been well described (Fig. 1E), whereas the regulation of Rac in follower cells remains unexplored.

To test for the requirement for Rac in follower cells, we expressed RacDN clonally in a varying number of border cells using the FLP-OUT technique. We reasoned that if every cell contributes to cluster movement, then as more cells within the cluster express RacDN we expected a concomitant decrease in cluster motility. So, we compared migration of clusters with different numbers of cells expressing UAS-moesinGFP and either a control (UAS-*white* RNAi) or UAS-RacDN. All clusters containing cells expressing *white* RNAi completed migration by stage 10, regardless of the number of labeled cells (Fig 1F). In contrast, the more RacDN-expressing cells in a cluster, the more severe the migration defect (Fig. 1F). None of the clusters with four cells expressing RacDN completed migration, and even a single RacDN-expressing cell could impede migration in 10% of clusters. We conclude that Rac activity in every migratory cell of the cluster contributes to their collective movement.

### Rac is required for follower cell crawling

While lead cell protrusions are obvious when the entire border cell cluster is labeled, follower cell morphology and dynamics are difficult to discern. Therefore we expressed MoesinGFP in subsets of border cells and made time lapse movies. Control clones expressing MoesinGFP and RNAi against the *white* gene showed that follower cells typically extend dynamic protrusions (Fig. 2A). Rather than protruding outward between nurse cells, as the lead cells do, followers actively crawl over other cells of the cluster (Movie 2). In contrast, RacDN-expressing cells exhibit little actin dynamics (Fig. 2B, Movie 3). In fixed imaging, we observed that labeled control follower cells typically exhibit a relatively compact morphology while extending lateral protrusions around E-cadherin boundaries with neighboring cells, as they crawl over and enwrap one another (Fig 2C-C”‘, Movies 2 and 4). In contrast, RacDN-expressing follower cells, which were most commonly in the rear position, exhibited an extremely elongated morphology such that the majority of the cell lacked contact with and was not integrated into the cluster (Fig 2D-D’, Movie 5). Thus follower cells require Rac for actin dynamics, which promote cell motility and cluster compactness.

**Figure 2:**
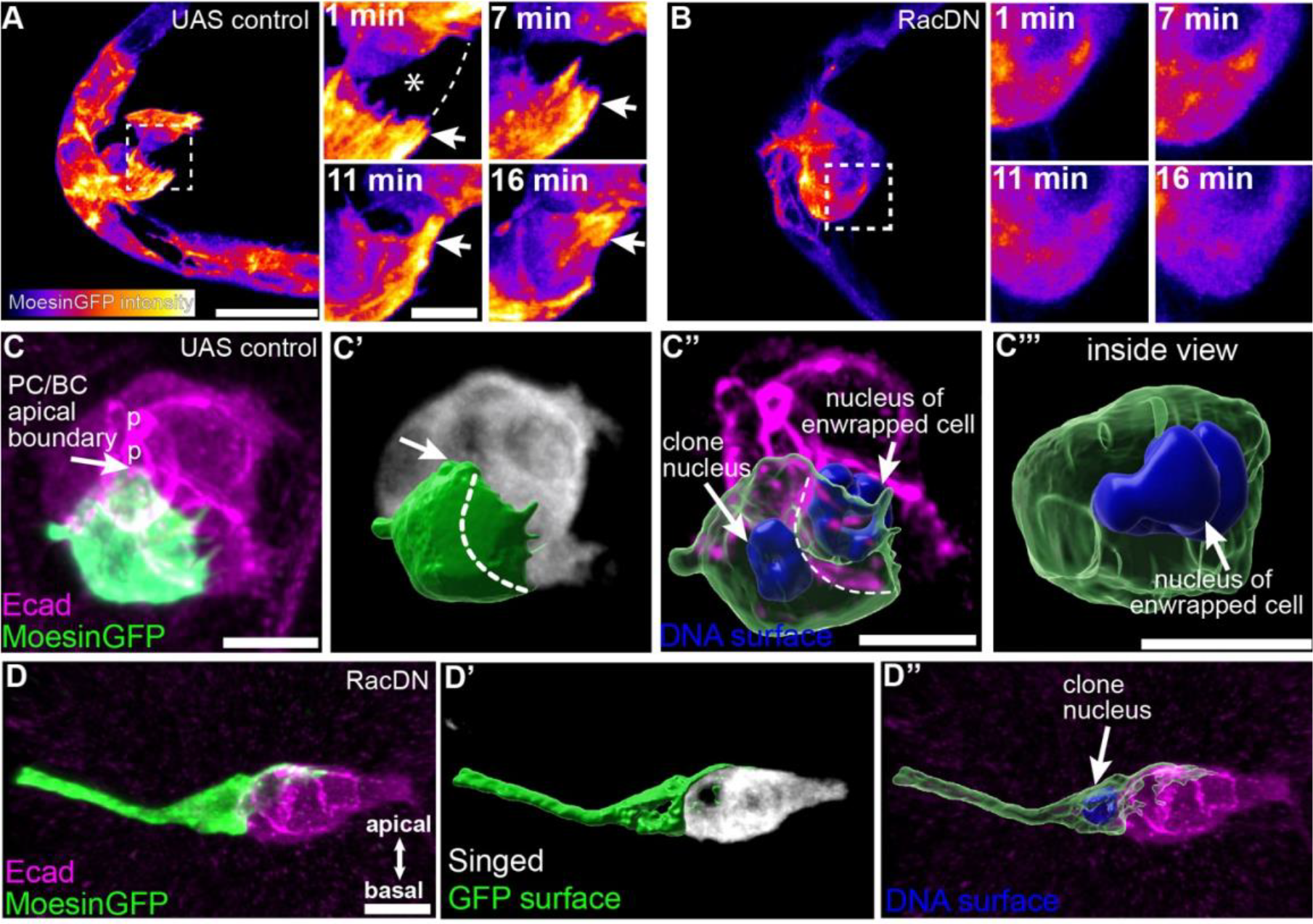
Rac is required for follower cell crawling. (A-D) Intensity color-coded live imaging of AyFLPout clones at time zero expressing UAS-moesinGFP and co-expressing either (A) UAS-*w*RNAi or (B) UAS-RacDN. Right, zooms of the boxed region. Arrows mark protrusions from crawling follower cells on an unmarked cell (*= unmarked cell, dashed line= approximate cell boundary). (C-D) Single border cell clones expressing UAS-moesinGFP and either (C) UAS-*w*RNAi or (D) UAS-RacDN. Ecad (magenta) and Singed (white). Polar cell apical surface marked with “p”. GFP surface reconstruction (green). (C’) Dashed line marks the junction between the clone and neighboring cell. (C’-C’’’ and D’-D’’) show same clone with surface overlay. (C’’’) Rotated view of the neighboring cell. All scale bars= 20 µm except C-C’’’ which are 10 µm.

### The basally localized RacGEF, Cdep, is required for follower cell crawling and cluster cohesion

The Rac GEFs sponge (Spg), myoblast city (Mbc) and Vav are required in the lead cell for protrusion (Bianco et al., 2007; Fernández-Espartero et al., 2013; Wang et al., 2018). To determine which GEF or GEFs might be required in follower cells, we carried out an RNAi screen of 26 putative Rac GEFs and identified one, Cdep, that is required for the crawling behavior of follower border cells. In contrast to control clusters, which move as a compact and cohesive unit (Fig. 3A, Movie 6), Cdep RNAi-expressing border cell clusters do not (Fig. 3B and Fig. S1, Movie 7). Two different RNAi lines effectively knocked down Cdep protein (Fig. S1A-B’) and caused similar defects. *Cdep* is the predicted *Drosophila* ortholog of human *FARP1* (29% identical and 42% similar) and *FARP2* (27% identical and 40% similar).

**Figure 3:**
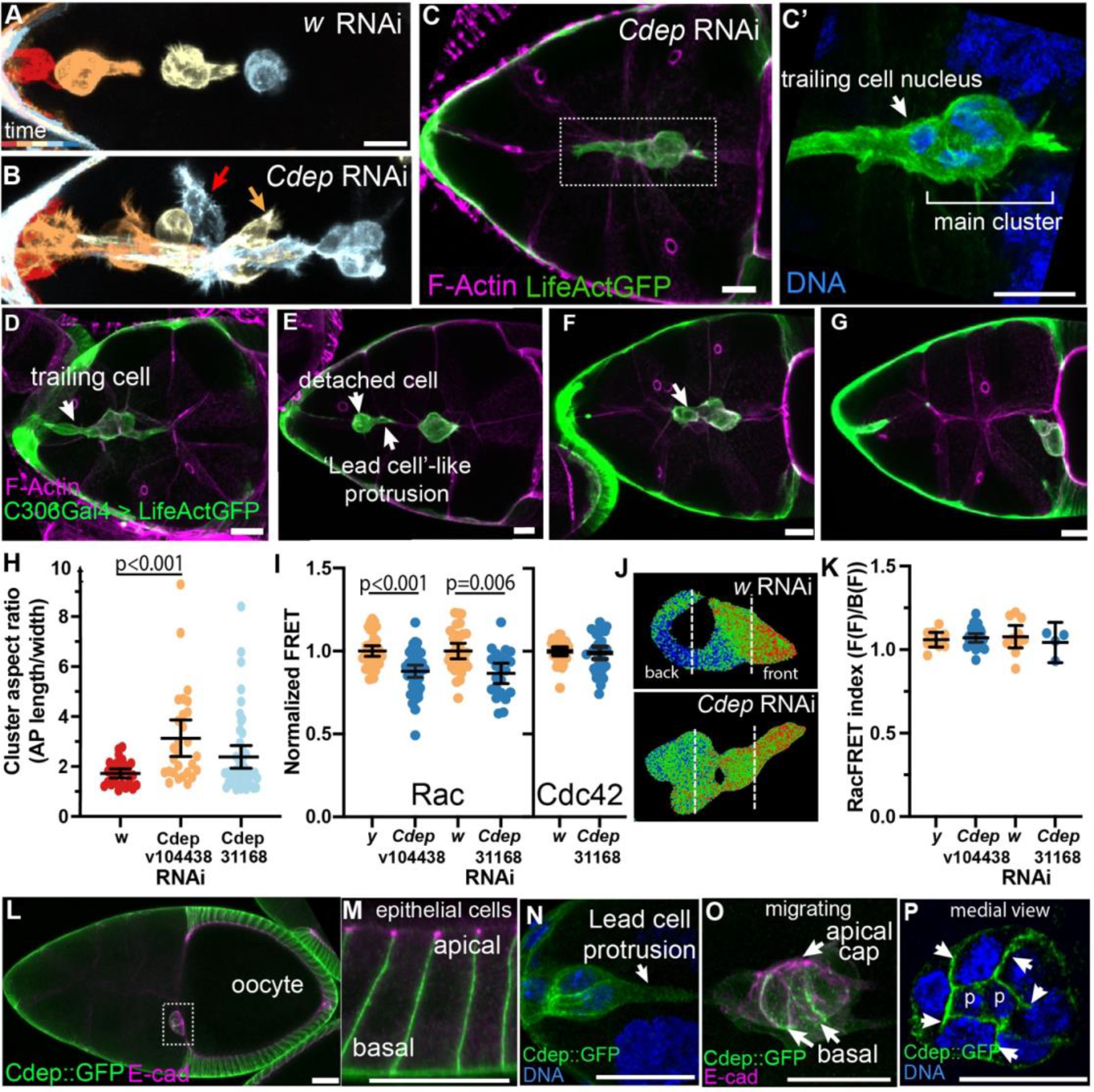
The basally localized RacGEF, Cdep, is required for follower cell crawling behavior and cluster cohesion. (A-B) 30 min interval color coded images from 3.5 hr time lapse movies for RNAi indicated. Red arrow indicates a detached border cell, and orange arrow indicates ectopic protrusion. (C-G) Representative projections of border cell clusters expressing UAS-*Cdep*RNAi stained for F-actin (magenta) and slbo>LifeActGFP (green). (C-D) trailing cells, (E) single cells falling off the cluster, (F-G) loss of cohesion. (H) Dot plot of cluster aspect ratio. (I) Analysis of c306Gal4, UAS-RacFRET or UAS-Cdc42FRET intensities and co-expressing control UAS-*w*RNAi/UAS-*y*RNAi or UAS-*Cdep*RNAi. Each dot is one cluster. (J) Representative images of RacFRET. (K) Front/back RacFRET intensity ratio of protrusive clusters in (I). (L) Projection of stage 10 egg chamber stained for Cdep::GFP (green) and Ecadherin (magenta). Dashed box indicates docked border cell cluster. (M) Zoom of epithelial follicle cells. (N) Delaminating border cells stained for Cdep (green). (O) Migrating border cell cluster stained as in (L). (P) Medial view of a border cell cluster with arrows marking border/border cell boundaries and “p” marking polar cell nuclei. All scale bars= 20 µm. Statistical significance tested using one-way ANOVA with Kruskal-Wallace (H) and Tukey (I and K) post hoc analysis.

Wild type border cells initiate migration with a single lead cell protrusion, and when individual frames taken at ∼30 minute intervals from a time lapse movie are displayed in different colors in a single figure, the spatial separation between border cell clusters at different time points reflects cohesive and coordinated cluster movement (Fig. 3A). The crawling defect in Cdep knockdown clusters (Movie 7) causes abnormally elongated clusters (Fig. 3B and Fig. S1D-L). Similar to RacDN clones, some cells of the cluster do not crawl forward; however, they have dynamic filopodia not observed in RacDN-expressing cells (Movie 7). The white color in Fig. 3B indicates failure to move and thus overlap of multiple time points. Cluster cohesion was reduced and some cells protruded ectopically (Fig. 3B, arrows).

In fixed imaging, we also observed trailing cells and split clusters consistent with an inability of the follower cells to crawl on one another to maintain cluster cohesion (Fig. 3C-G). In some cases a single cell showed a trailing phenotype similar to clonal expression of RacDN in a single cell (Fig. 3C, C’), while in other cases a single cell detached from the cluster (Fig. 3E). Interestingly, such cells could make a lead-cell-like protrusion (Fig. 3E), consistent with Cdep being required for follower cell behavior. The trailing phenotype and separation of follower cells caused an increase in the cluster aspect ratio in Cdep knockdown clusters (Fig. 3H).

To determine whether *Cdep* is a RacGEF, we expressed *Cdep* or control RNAi for the *yellow* (*y*) or *w* gene in combination with a validated RacFRET activity probe (Wang et al., 2010). Indeed, knockdown of *Cdep* with two independent RNAi constructs significantly lowered RacFRET but not Cdc42FRET compared to the controls, indicating *Cdep* is a *bona fide* RacGEF in the border cells (Fig. 3I). Interestingly we found that while overall levels of RacFRET were reduced, the front-back bias of Rac activity was unchanged in Cdep knockdown (Fig. 3J and K), supporting our hypothesis that Cdep functions predominantly outside of the lead cell protrusion.

To gain further insight into Cdep function, we examined its localization. We found that an endogenously GFP-tagged Cdep (Cdep::GFP) is expressed in all follicle cells, including border cells (Fig. 3L-P). Co-staining with the cell adhesion molecule E-cadherin, which is enriched in subapical adherens junctions, showed that Cdep localizes to basolateral membranes in the follicular epithelium (Fig. 3M) and in border cell clusters (Fig 3N-P). Cdep::GFP is enriched at cell-cell contacts but not in lead cell protrusions (3N). High resolution imaging of migrating clusters showed that Cdep is particularly enriched basally at polar/border cell contacts (Fig. 3O) and at border/border cell membranes (Fig. 3N-P).

### Apicobasal polarity proteins regulate Cdep localization

Epithelial follicle cells exhibit classical apicobasal polarity with apically localized atypical protein kinase C (aPKC), lateral Scrib, and a basal basement membrane (Fig 4A and B). Border cells retain some apicobasal polarity as they migrate (Pinheiro and Montell, 2004; Godt and Tepass, 2009). However, polarity is clearest in contacts between polar cells and between border cells and polar cells (Fig. 4C). As border cells delaminate, they undergo a 90° turn, which causes the apical domain to orient roughly perpendicular to the direction of migration (Fig. 4D-F”). At the end of migration, they turn again to dock their apical surfaces to the oocyte (Fig. 4G-G”). The basolateral protein Scrib is found on basal and lateral membranes between border cells and polar cells and the apical polarity determinant, aPKC, is enriched in apical junctions (Fig. 4C). Crumbs, Par, and Scrib modules are required for border cell migration (Li et al., 2011; Pinheiro and Montell, 2004; Szafranski and Goode, 2004; Wang et al., 2018; Zhao et al., 2008), but the mechanism by which they act is not yet clear.

**Figure 4:**
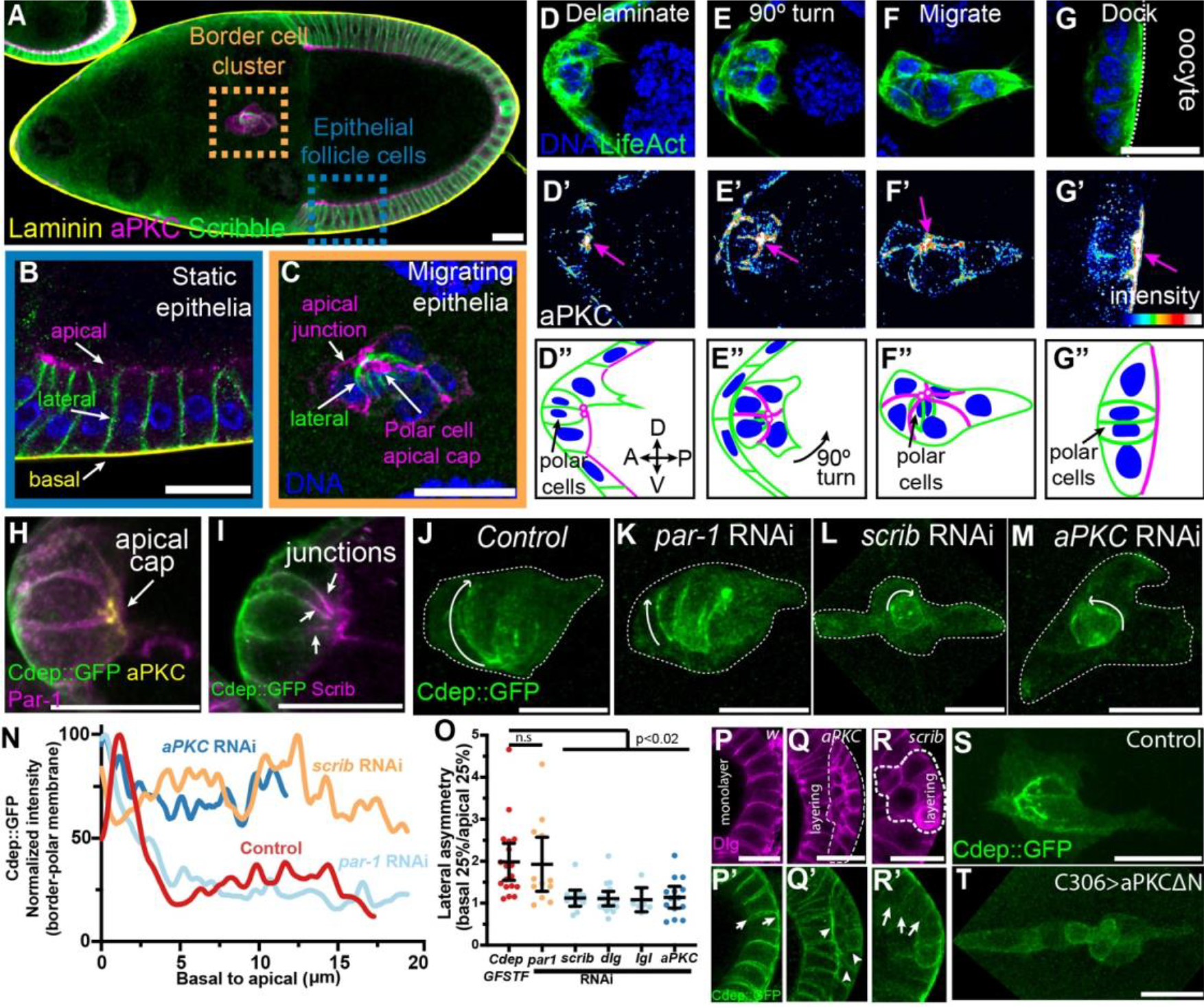
Core polarity modules localize Cdep in border cell clusters. (A) Projected stage 9 egg chamber with migrating border cells stained for Laminin with peanut lectin (yellow), aPKC on apical membrane (magenta), and Scribble on lateral membrane (green). (B) Epithelial follicle cell and (C) border cell zooms of (A) with DNA (blue). (D-G’’) Stages of border cell migration from stage 9-10 egg chambers with DNA (blue), (D-G) Actin staining from *slbo*>LifeActinGFP reporter (green), (D’-G’) aPKC staining (color coded by intensity) with arrows to denote the apical cap of the polar cells, and (D’’-G’’) schematic representations of D-G’. Axis in D’’ indicates anterior, A, posterior, P, dorsal, D, and ventral, V. (H) Overlay of a border cell cluster stained for endogenous Cdep::GFP (green), Par-1 (magenta), and aPKC (yellow). (I) Overlay of a border cell cluster stained for Cdep::GFP (green), and Scribble (magenta). (J-M) Representative images of border cell clusters expressing Cdep::GFP in combination with indicated RNAi. Arrow points toward apical. (N) Representative line scans along the polar/border cell boundary from basal to apical for genotypes in (J-M). (O) Dot plot of basal to apical asymmetry of Cdep at polar/border cell membrane for indicated knockdowns. Each dot is an individual border cell cluster. Statistical significance was tested using one-way ANOVA with Kruskal-Wallace post hoc. (P-R’) Representative images of Dlg (magenta) and Cdep::GFP (green) in control (P,P’), *aPKC* RNAi (Q-Q’) and *scrib* RNAi (R-R’). Layering is outlined. (S) Control and (T) UAS-aPKCΔN expressing border cell clusters during migration. All scale bars= 20 µm.

To interrogate the relationship between Cdep and the polarity modules, we compared Cdep localization to that of aPKC, the Scrib module, and the basal kinase Par-1. Cdep::GFP membrane localization partially overlapped with Par-1 (Fig. 4H, Fig. S2A-A’’’), although Cdep was concentrated even more basally than Par-1 in these cells. Cdep was undetectable in apical membranes and thus did not colocalize with aPKC at the polar cell/border cell boundary. Cdep overlaps with Scribble in lateral membranes though Cdep is concentrated more basally, while Scrib is most enriched just basal to the adherens junction (Fig. 4I and Fig. S2B-B”‘).

To test if Cdep membrane localization and polarization depends on apicobasal polarity proteins, we examined Cdep::GFP in cells expressing RNAi lines against *Par-1, aPKC*, and *scrib*. Par-1 knockdown did not detectably change the localization of GFP::Cdep (Fig. 4K). In contrast, knockdown of either Scrib or aPKC caused a general reduction in Cdep::GFP membrane association and a more diffuse cytoplasmic signal (Fig. 4L and M). Line scans of border cell-polar cell contacts from basal to apical showed that Par-1 knockdown did not change Cdep asymmetry whereas Scrib and aPKC did (Fig. 4N and O). Dlg and Lgl knockdowns phenocopied Scrib (Fig. 4O).

Analysis of posterior epithelial follicle cells from the same egg chambers also showed a difference in Cdep membrane localization.

Knockdown of either Scrib or aPKC causes multiple layers of follicle cells to form, characteristic of epithelial polarity defects (Fig. 4P-R). Whereas Cdep::GFP is normally excluded from apical domains (Fig. 4P, P’), Cdep was present on all cell surfaces within the layering event and was enriched between layers (Fig. 4Q, Q’). Conversely, knockdown of Scrib led to reduced membrane accumulation of Cdep::GFP on lateral membranes and more diffuse cytoplasmic signal (Fig. 4R-R’). The data from both border cells and posterior follicle cells suggest that the Scrib module promotes Cdep membrane association whereas aPKC antagonizes it.

A prediction then is that expression of a constitutively active form of aPKC would reduce Cdep membrane association. As expected, when we overexpressed a constitutively active form of aPKC lacking its NH_2_-terminus (aPKCΔN) (Betschinger et al., 2003), Cdep::GFP failed to associate with basal membranes and appeared more diffusely cytoplasmic in both border cells (Fig. 4S and T) and epithelial cells (Fig S2C and D). We conclude that the Scrib module promotes membrane association while aPKC excludes Cdep from the apical domain.

### The Scrib module promotes cluster integrity independent of tumor suppression

To investigate the mechanism by which Scrib, Dlg, and Lgl regulate border cell migration, we used validated RNAi lines to knock down their expression. However these proteins are tumor suppressors, and mutations cause hyperproliferation and multilayering of follicle cells that at best complicate and at worst preclude analysis of border cell migration (Goode and Perrimon, 1997; Goode et al., 2005). To circumvent this issue, we conditionally expressed RNAi lines targeting *scrib, dlg* or *lgl* using c306Gal4 (Manseau et al., 1997) and the temperature-sensitive repressor tubGal80^ts^. RNAi against the *white* gene was used as a control. Incubating c306Gal4, *tub*Gal80^ts^, UAS-RNAi flies at 30°C (to inactivate Gal80^ts^) for 72 hours was long enough to achieve knockdown of Scrib, Dlg and Lgl (Fig. S3A-F) without causing multilayering or proliferative tumors. LifeActGFP labeled the border cell F-actin cytoskeleton and showed that, in contrast to controls (Fig. 5A), Scrib, Dlg or Lgl knockdown caused cluster elongation (Fig. 5B-D), individual cells splitting off (Fig. 5B, C, red arrows), and ectopic protrusions from follower cells (Fig. 5B-D, orange arrows). Compared to control clusters, knockdown of the Scrib module reduced the percentage of border cells that complete migration (Fig. 5E-F), and increased overall cluster aspect ratios by ∼50% (Fig. 5G). Multiple RNAi lines against each gene caused similar effects (Fig. 5F). Longer incubation times resulted in multilayering, as expected (Fig. S3G-I”).

**Figure 5.**
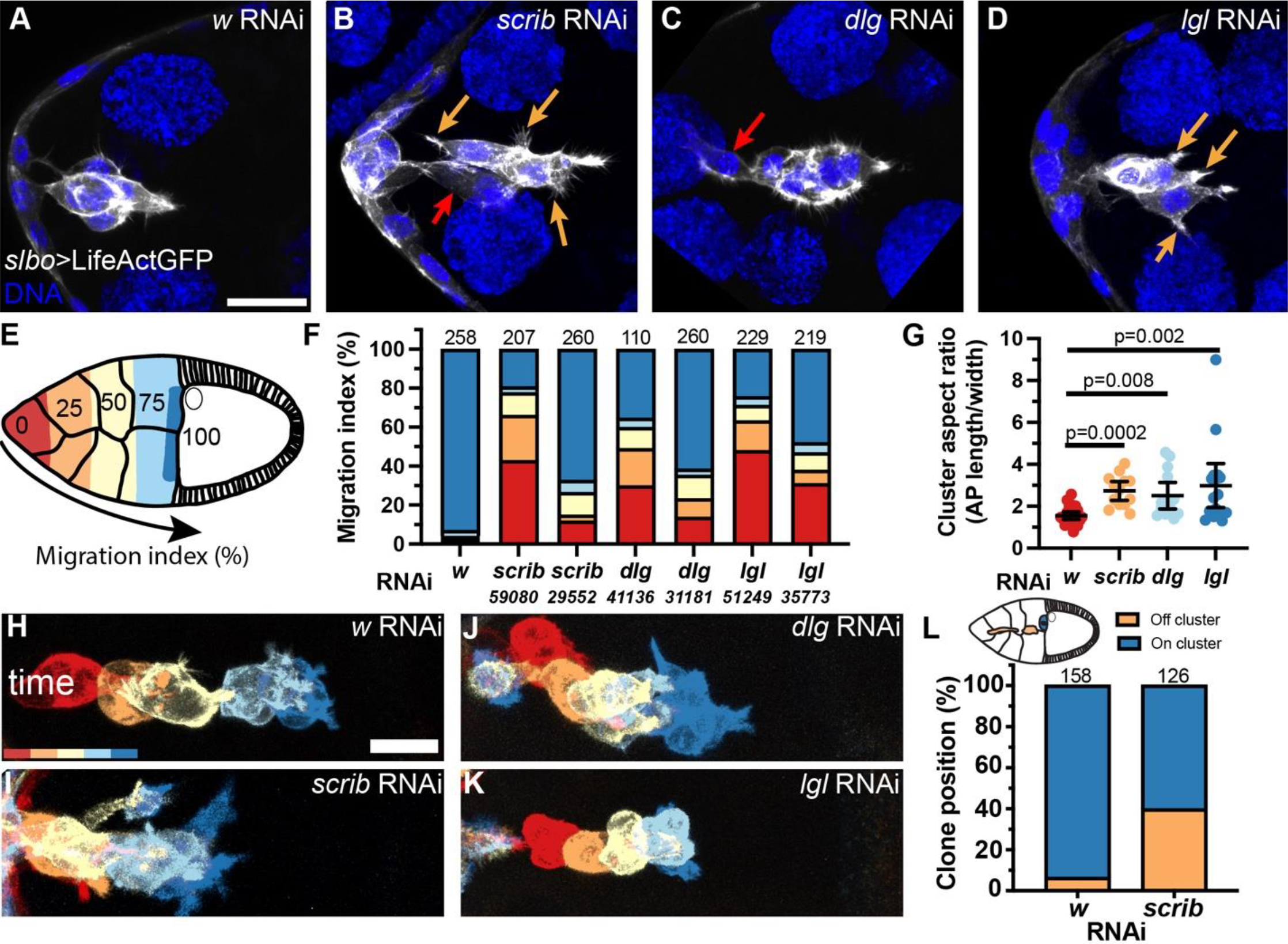
The Scrib module coordinates cluster cohesion, detachment, and migration dynamics. (A-D) Representative projections of migrating border cell clusters expressing c306Gal4, *slbo*>LifeActGFP, *tub*Gal80^ts^, and indicated UAS-RNAi. Lifeact (white) and DNA (blue). Cells with ectopic protrusions or that fell off the cluster marked with orange or red arrows, respectively. (E) Color coded schematic of migration index measurements in stage 10 egg chambers. (F) Frequency distribution of border cell migration at stage 10. (G) Dot plot of cluster aspect ratio. (H-K) 30 min color coded images from 4 hr time lapse movies for RNAi indicated. (L) Schematic and proportion of stage 10 egg chambers with clonal border cells on or off the cluster expressing the indicated UAS-RNAi under the control of HsFlp>AyGal4. All scale bars= 20µm. Statistical significance tested using one-way ANOVA with Kruskal-Wallace post hoc analysis (G).

Live imaging confirmed that clusters progressively elongate throughout their migration (Fig. 5H-K, Fig. S3J-Q, Movies 8 and 9) similar to Cdep knockdown. In the most extreme cases of cluster cohesion loss, individual cells separated from the cluster. To quantify how frequently this occurred we clonally knocked down Scrib in cells of the cluster and observed that ∼40% of *scrib* RNAi containing clusters had cells that completely detached from their neighboring wild type counterparts compared to ∼5% in *white* RNAi (Fig 5L). These results indicate that Scrib knockdown reduces cluster cohesion and impedes cluster migration.

### Localizing Cdep to basal membranes partially suppresses Scrib phenotypes

Scrib knockdown causes diverse phenotypic effects and Scrib interacts with dozens of binding partners (Bonello and Peifer, 2019), raising the question as to which effectors mediate Scrib function. Since Scrib knockdown disrupts Cdep membrane localization, we wondered which, if any, Scrib functions might be mediated by Cdep. So we used the GrabFP system to re-localize Cdep::GFP to basal membranes in Scrib knockdown cells. GrabFP is an anti-GFP nanobody-based trap that was developed to concentrate GFP tagged proteins in specific subcellular locations (Harmansa et al., 2017). We used GrabFP-Basal, in which Nrv1 is fused to the anti-GFP nanobody to target it to basolateral membranes. As a negative control we used ExGrabFP-Basal, in which the anti-GFP nanobody is expressed on the exterior of the cell and so should not interact with intracellular Cdep. We expressed either GrabFP-Basal or ExGrabFP-Basal with Cdep::GFP and UAS-*scrib* RNAi with c306Gal4 in the absence of tubGal80^ts^. Without tubGal80^ts^ to attenuate the Scrib knockdown phenotype, egg chambers had extensive anterior follicle cell layering (Fig 6A and D, white arrows) in addition to border cell migration phenotypes (Fig 6B and E) and elongation of the border cell cluster (Fig 6C and F). Under these conditions, most egg chambers eventually became necrotic (Fig. 6D), hampering our ability to quantify border cell motility. However, expression of GrabFP-Basal efficiently restored basolateral localization of Cdep::GFP in *scrib* RNAi-expressing follicle cells (Fig. 6A-C). Basolateral localization of Cdep::GFP also ameliorated multiple Scrib knockdown phenotypes including egg chamber death (Fig 6A and D) and follicle cell layering frequency and severity (Fig. 6G and H) compared to the ExGrabFP-Basal control. In the healthy stage 10 egg chambers, we observed a significant reduction in the number of anterior layering events with GrabFP-Basal vs ExGrabFP-Basal (Fig. 6I). Cluster cohesion was also significantly improved in GrabFP-Basal vs ExGrabFP-Basal (Fig. 6J). We conclude that basolateral membrane targeting of Cdep is sufficient to improve egg chamber viability, restore follicle cell epithelial polarity, and promote border cell cluster cohesion.

**Figure 6:**
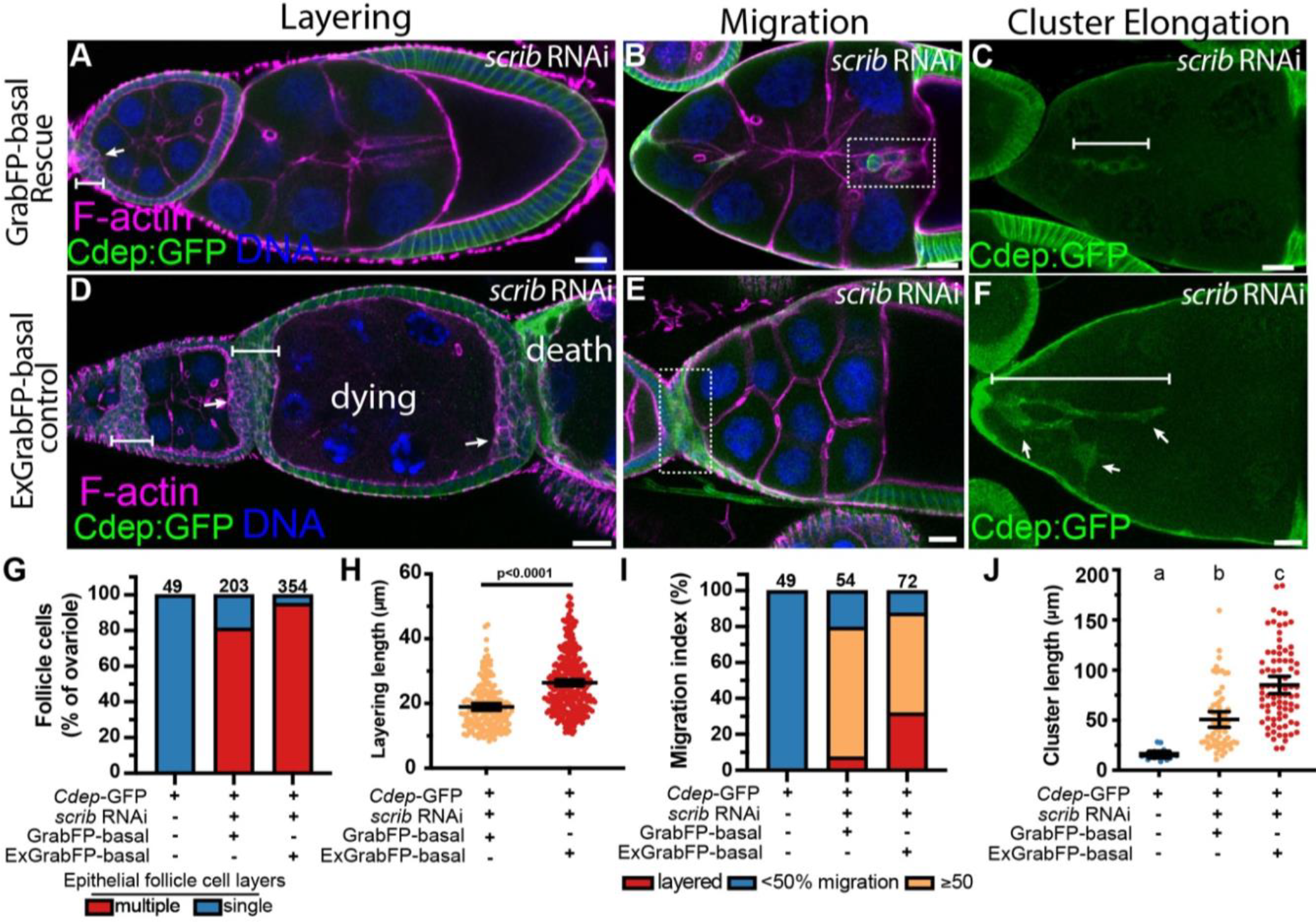
Re-localizing Cdep to the basal membrane suppresses border cell epithelial layering and cluster elongation caused by Scrib module knockdown. (A-F) Egg chambers expressing (A-C) UAS-GrabFP-basal or (D-F) UAS-ExGrabFP-basal and co-expressing UAS-Scrib RNAi, stained for F-actin (magenta) and DNA (blue). (A and D) Layering throughout the egg chamber (arrow). Bars represent the length measurement for anterior layering. (B and E) Dotted box indicates the border cell cluster. (C and F) Bar represents the length measurement for cluster elongation. (G) Quantification of ovariale strands where either no layering was observed (blue), or any layering (orange) anywhere within a single ovariole strand. (H) Quantification of anterior layering size (I) Frequency distribution of migration defects in stage 10 egg chambers. (J) Quantification of cluster elongation as the length of the longest line that can be drawn through the anterior-posterior axis of the border cell cluster. Statistical significance was tested using one-way ANOVA with Kruskal-Wallace post hoc analysis. All scale bars= 20 µm.

In further support of the sufficiency of basolateral Cdep to rescue epithelial follicle cell defects, we found that basolaterally localized Cdep also reduced the frequency of ovariole strands containing even a single egg chamber of any stage with any epithelial layering (Fig. 6G). In addition, even in those egg chambers that did have multiple layers, basolateral re-localization of Cdep significantly reduced the number of cell layers (Fig. 6H) (p<.0001). These findings suggest that a major function of the Scrib module during egg chamber development is to localize Cdep to basolateral membranes.

## Discussion

### Follower cell crawling depends on Rac and promotes cluster cohesion

Collective cell migration is a widespread phenomenon that drives much of embryonic development, wound healing, and tumor metastasis. In migrating collectives, there are typically leader cells that extend protrusions outward from the group to steer it (Haney et al., 2018; Khalil and de Rooij, 2019; Theveneau and Linker, 2017; Zhang et al., 2019). The functions of follower cells are less clear (Qin et al., 2021), especially *in vivo*. Follower cell contributions have primarily been studied in collective sheet migration *in vitro*, rather than in small clusters. But clusters are emerging as major contributors to metastasis (Aceto et al., 2014; Au et al., 2016; Cheung and Ewald, 2016; Cheung et al., 2016; Conod et al., 2022), so it is of interest to understand the mechanisms by which clusters maintain cohesion as they move.

Cohesion of collectively migrating cell sheets has been primarily attributed to cadherin-mediated adhesion between cells (Khalil and de Rooij, 2019). In border cells, as in collective sheet migration, E-cadherin mechanically couples leaders to followers, inhibiting followers from protruding outward between nurse cells (Cai et al., 2014). However, knockdown of E-cadherin in outer border cells does not cause a loss of cluster cohesion; rather the clusters lack a leader and crawl and tumble in random directions (Cai et al., 2014).

Based on the data presented here, we propose that border cell cluster cohesion is mediated at least in part by follower cells enwrapping one another as they crawl (Fig. 7). In many ways the E-cadherin knockdown phenotype and the Scrib/Cdep knockdown phenotypes are complementary to one another. In E-cadherin knockdowns, cluster cohesion is maintained and follower cell crawling is active, while lead cell protrusions are lost (Fig 7 of this work and Cai et al., 2014, Figure 1F, Movie S2). Conversely, in Scrib or Cdep knockdowns, leading protrusions are present, but follower cells are impaired in motility, so that the cluster becomes elongated, and some cells completely detach. In contrast, in sheets of cells migrating on matrix, basal cryptic protrusions mediate cell-substrate interactions and traction forces transmit tension to E-cadherin-mediated cell-cell contacts to coordinate the collective motion (Ng et al., 2012). Thus the mechanism for maintaining cohesion and coordinated movement of small clusters of cells squeezing between other cells may differ substantially from the mechanism in a sheet of cells moving on matrix, even though both types of collectives contain leaders and followers.

**Figure 7:**
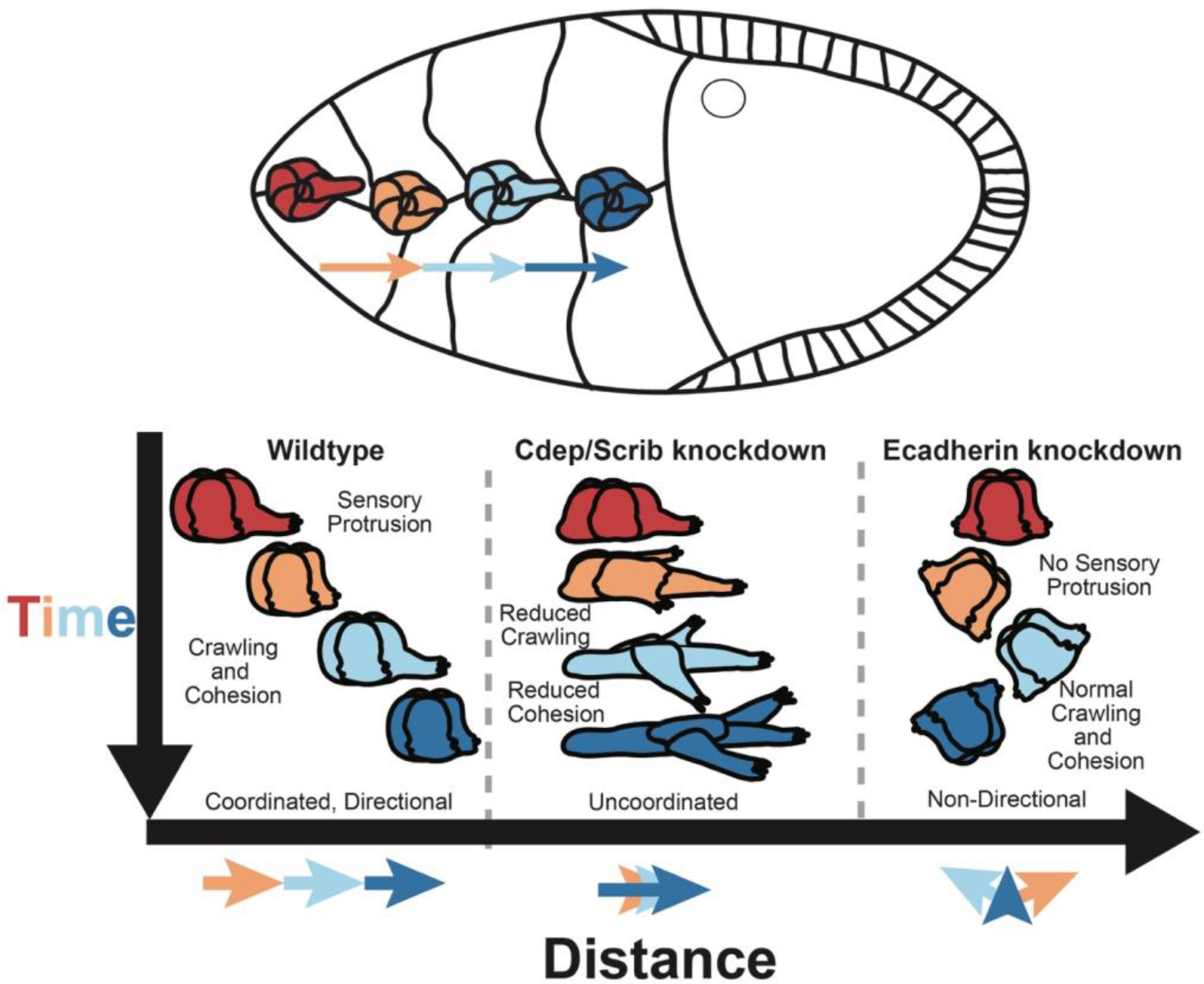
An integrated view of collective border cell migration. Illustration of wild type border cell cluster migration.olor indicates sequential time points. Comparison of features of normal border cell leader and follower cell behavior and cluster movement to those of Cdep or Scrib knockdown and the complementary defects caused by E-cadherin knockdown (based on Cai et al., 2014, Movie S2). In wild type, the front cell extends a large protrusion that senses chemoattractants (Duchek and Rørth, 2000; Duchek et al., 2001; McDonald et al., 2004) and small spaces (Dai et al., 2020) and is shaped and stabilized by E-cadherin-mediated adhesion (Cai et al., 2014) and steers the cluster. Cluster cohesion and movement also require basal, Rac-dependent crawling in follower cells. Knockdown of Cdep or Scrib disrupts crawling and thus cluster cohesion and coordinated movement. E-cadherin knockdown causes loss of lead cell protrusion and directional migration but cells within the cluster continue to crawl, maintaining cohesion and leading to random movement.

In border cells, the lead cell protrusion was initially proposed to function as a grapple that would pull the rest of the cluster forward (Fulga and Rørth, 2002; Rørth, 2009). This idea implied that the follower cells might be passive passengers, thus much work focused on elucidating the molecular pathways regulating Rac in the lead cell protrusion. However the grapple model was based on fixed tissue imaging.

Live imaging has revealed that 90% of lead protrusions retract and that cluster speed is uncorrelated with extension or retraction of protrusions (Mishra et al., 2019b). These observations lead to a model in which the large forward-directed protrusion is primarily a sensory structure, probing for chemoattractants, traction, and physical spaces (Dai et al., 2020), rather than a grapple.

If the lead protrusion does not function as a grapple, then how does the cluster actually move? Here we show that all cells contribute to cluster movement. We show that Rac is required in each cell and that the molecular pathway regulating Rac-mediated crawling differs from the regulators of lead cell protrusion. And whereas the Rac GEFs in the lead cell protrusion are activated downstream of chemoattractant receptor signaling, Cdep localization is regulated by the basolateral proteins Scrib, Dlg and Lgl rendering follower cells more dependent on polarity than chemoattractant signaling.

Scrib, Dlg and Lgl are best known as tumor suppressors (Bilder et al., 2000; Brumby and Richardson, 2003; Hariharan and Bilder, 2006), mutation of which causes tissue overgrowth. Scrib loss of function causes epithelial to mesenchymal transition and increased single cell invasion in cells expressing oncogenic Ras (Igaki et al., 2006; Pagliarini and Xu, 2003; Wu et al., 2010). Paradoxically though, Scrib, Dlg, and Lgl are infrequently mutated in human cancers (Lin et al., 2015; Santoni et al., 2020). The results here provide a possible explanation. We show that Scrib loss of function impedes collective motility of border cell clusters. If tumors spread collectively and Scrib is similarly required for cluster cohesion and coordinated migration, then the function identified here could provide a selective pressure against Scrib loss-of-function mutations.

Polarity signaling complexes and Rho GTPase activities appear to be well-conserved between border cells and mammalian cells that migrate collectively, and a basal Rac GEF dependent upon Scribble has even been predicted but not yet identified (Zegers and Friedl, 2014). The results presented here suggest that FARP1 and FARP2 would be excellent candidates to mediate cluster cohesion in collectively migrating mammalian cell clusters downstream of Scribble. Neither has been extensively studied, however the little that has been reported is intriguing. *FARP1* is required for migration of lung adenocarcinoma cells (Cooke et al., 2021), and *FARP2* promotes collective invasion of colorectal carcinoma (Libanje et al., 2019). Colorectal carcinomas invade and spread as small groups of cells with apicobasal polarity. It will be interesting to determine whether it serves a similar function in that context.

## Materials and Methods

### *Drosophila* husbandry and genetics

A detailed list of all fly strains used in this study, their source and the figures in which they were used is documented in Table 1. Strains outside of lab stocks were ordered from the Bloomington Drosophila Stock Center, the Vienna Drosophila Resource Center, or kindly donated (see table 1). Most crosses were grown at 25ºC and flies were fattened by yeast addition on an *ad libitum* basis for three days at 29ºC before dissection. The exceptions were for crosses containing the conditional expression construct, tubGal80ts, in which flies were grown at 18ºC and transitioned to 30ºC for fattening. tubGal80ts is a temperature dependent repressor of Gal4 activity, thus at restrictive temperature (18ºC) Gal4 remains inactive and at permissive temperature (30ºC), Gal80ts is inactive therefore allowing Gal4 to be active. Experiments that required clonal border cells expressing RNAi and a fluorescent clonal marker were generated using the FLP/FRT clonal marking system. RNAi stocks were crossed to either hsp70-FLP; Acty17bGal4, UAS-moesin-GFP or hsp70-FLP; Tub[FRT]CD2[FRT]Gal4, UAS-GFP/SM5a were heat-shocked two times at 37ºC with at least 6 hours between each heat shock. Flies were fattened for 48 hr and dissected as stated below for fixed staining or live imaging.

### Immunohistochemistry

5-7 fattened adult female flies were dissected in Schneider’s Medium (ThermoFisher, catalog #21720001) supplemented with 20% fetal bovine serum and 1x antimycotic/antibiotic (VWR, catalog #45000-616). Ovarioles were separated from the muscle sheath using Dumont style 5 stainless steel forceps (Electron Microscopy Sciences) before fixation. Egg chambers were fixed in 4% paraformaldehyde in phosphate buffered saline (PBS) for a total of 15 min at room temperature before washing three times 15 min with PBS+0.2% TritonX-100 (PBST). Primary antibodies were diluted in PBST and ovarioles were incubated overnight at 4ºC. Egg chambers were then washed with PBST three times at room temperature, incubated in fluorescently conjugated secondary antibodies (Invitrogen, all AlexaFluor conjugated goat antibodies) in PBST for 2 hr, and washed three times more with PBST. Samples were then mounted in VectaShield (Vector Laboratories, catalog #H-1000) mounting medium and stored at 4ºC until imaging. Egg chambers were mounted on a #1.5 coverglass and imaged with either a Zeiss LSM780 or LSM800 confocal microscope using either a PlanAPO 20x 1.2NA or a PlanAPO 40x 1.4NA objective. All image settings were kept exactly the same within each experiment.

Antibodies used in this study include: rabbit anti-Scrib (kind gift from Dr. Chris Doe, 1:2000), guinea pig anti-Scrib (kind gift from Dr. David Bilder, 1:500), Rabbit anti-Par-1 (a kind gift from Dr. Jocelyn McDonald, 1:500), mouse anti-PKCζ (sc-216) and rabbit anti-PKCζ (sc-17781), mouse anti-HA (sc-7392, 1:1000), and rabbit anti-LGL (sc-9826) were from Santa Cruz Biotechnology and used at 1:500, mouse anti-Dlg (4F3, 1:25), mouse anti-Cut (2B10, 1:100) and rat anti-Ecad (DEcad, 1:20) were from the Developmental Studies Hybridoma Bank, rabbit anti-GFP (A11122, 1:1000) from ThermoFisher, chicken anti-GFP (ab13970, 1:1000) from Abcam. Lectin PNA Alexa 647 was purchased from Thermo Fisher Scientific (L32460).

### Live imaging

Egg chambers were dissected from 5-7 fattened adult female flies in Schneider’s Medium supplemented with 20% fetal bovine serum and 1x antimycotic/antibiotic. Egg chambers were washed two times with dissecting medium and mounted on Lumox imaging dishes in medium containing 0.4 mg/mL bovine insulin and 1% low melt agarose. Egg chambers were imaged by confocal as above.

### Generating UAS-HA-Cdep

The RNeasy kit (Qiagen, catalog #74124) was used to isolate total RNA from the ovaries of six fattened female flies. RNA was converted to cDNA using the SuperScriptIII-First strand synthesis kit (ThermoFisher, catalog #18080-51). The following primers with overhangs for subsequent Infusion cloning into the pJFRC7 vector and encoding a N-terminal 2xHA tagg were used to amplify full length *Cdep* isoforms E/F. Forward:GCGGCCGCGGCTCGAGATGTACCCATACGATGTTCCAGATTACGCTGGATATCCATATGATGTTCCAGATTATGCTGGAATGTCCCTGGCCGACATGGG. Reverse:CATGCTGTGTTGGGCAACTAATCTAGAGGATCTTTGT.

Amplicons of tagged-*Cdep* were isolated by gel extraction after PCR amplification using Phusion high fidelity polymerase (New England Biolabs, catalog #E0553L). The amplicons were then cloned into the pJFRC7 5x UAS vector (ref) using Infusion cloning kit (Takara Bio, catalog #638948). Sequenced clones were sent to BestGene for injection and stock generation.

### Image processing and quantification

Quantification of apical-basal polarization of proteins along the border cell-polar cell boundary (Fig 1H/I, Fig 3D/E): When possible the apical cap of the polar cells was identified by either E-cadherin, Phalloidin/F-actin or aPKC staining. Using ImageJ a line scan with a thickness of .75 μm was drawn tracing the border cell-polar cell boundary using the F-actin channel as a guide. Measurements were then taken in the Cdep channel and averaged over 0.50 μm to smooth the signal and then normalized to the peak value within an individual border cell cluster. Representative line scans were chosen for the line graphs. To compare population differences we took the average of the 25% most basal signal divided by the 25% most apical signal for each sample. This ratiometric measurement of basal asymmetry was used for statistics and population level plotting. In a subset of polarity complex knockdown samples the apical cap could not be easily distinguished and an arbitrary pole was chosen as basal. In this condition, there was no bias seen in Cdep asymmetry regardless of orientation or start point chosen.

Quantification cluster aspect ratio (Fig 2G): A bounding box was manually drawn in FIJI around the border cell cluster in an anterior-posterior orientation. The aspect ratio of the bounding rectangle was then measured and plotted for each condition.

Quantification of cluster speed (Fig 2L): The center of mass of the border cell cluster was manually identified in each time point. The MTrackJ plugin was then used to calculate the velocity of the border cell cluster.

Quantification of Rac FRET (Fig 3D): FRET images were obtained during live cell imaging on a Zeiss LSM 780. A 458-nm laser was used for excitation of the sample. CFP and YFP images were collected simultaneously using channel 1 (464–502 nm) and channel 2 (517–570 nm) under a 40×/1.1 numerical aperture (NA) water immersion objective LD C-Apo lens. Images were taken at 16-bits with a frame size of 512 × 512 in the center of mass of the border cell cluster. CFP and YFP images were then processed using Fiji image analysis software as described before (Kardash ref). The final FRET image of the YFP/CFP ratio image was generated in ImageJ. The average FRET signal of the entire cluster was measured. All samples were observed after detachment and prior to docking.

Quantification of layering (Fig 5D/E): Images were taken of whole ovariale strands at a low zoom-level on a 20x objective using a Zeiss LSM 800 or Zeiss LSM 780. Images were 30-36 microns in thickness with a z-step of 2.5-3 microns. Images were then manually quantified in ImageJ by observing any location within the follicular epithelium that showed layering in any stage egg chamber. Quantification of layering was done only on anterior layering events in stage eight and nine egg chambers. The distance from the anterior tip of the egg chamber to the posterior-most cell within the layering event was measured to quantify the size of the layering event.

Quantification of cryptic protrusions (Fig6 D/E): Filopodia were manually marked at the distal tip and at where the protrusion meets the circumference of the border cell cluster. The distance between the XY coordinates at the tip vs base was then quantified with the distance formula: distance = sqrt(X_2_ - X_1_)^2^ + (Y_2_ - Y_1_)^2^).

### Graphing and statistical analyses

All data were analyzed in Excel (Microsoft). Graphs and statistics were generated in Prism 9 (GraphPad). For statistical analysis by one-way ANOVA, Bartlett’s Test was run to determine if significant differences in sample deviations warranted non-parametric analysis. In cases where deviations were not significantly different, parametric ANOVA was performed with Tukey post-hoc tests of significance. In cases where deviations were found to be significantly different, non-parametric ANOVA was run with Kruskal-Wallace post-hoc tests of significance.

## Supporting information

Maximum intensity projection of a control border cell cluster undergoing delamination.

Maximum intensity projection of control border cell clones expressing UAS-white RNAi and UAS-moesin::GFP

Maximum intensity projection of border cell clones expressing UAS-RacDN and UAS- moesin::GFP

Imaris reconstruction of a fixed border cell cluster with a single cell expressing UAS- white RNAi and UAS-moesin::GFP (green). Ecad in magenta.

Imaris reconstruction of a fixed control border cell cluster with a single cell expressing UAS-RacDN and UAS-moesin::GFP. Ecad in magenta

Maximum intensity projection of a control border cell cluster expressing UAS-white RNAi and UAS-LifeActGFP under the control of C306-Gal4

Maximum intensity projection of a border cell cluster expressing UAS-Cdep RNAi and UAS-LifeActGFP under the control of C306-Gal4.

Maximum intensity projection of a control border cell cluster expressing UAS-white RNAi under the control of C306Gal4

Maximum intensity projection of a border cell cluster expressing UAS-scrib RNAi under the control of C306Gal4

## Author Contributions

Conceptualization, J.P.C., J.A.M, and D.J.M.; Methodology J.P.C., J.A.M, and D.J.M.; Validation, J.P.C. and J.A.M; Formal Analysis, J.P.C. and J.A.M; Investigation, J.P.C. and J.A.M; Resources, J.P.C., J.A.M, and D.J.M.; Data Curation, J.P.C. and J.A.M; Writing - Original Draft, J.P.C., J.A.M, and D.J.M.; Writing - Review & Editing, J.P.C., J.A.M, and D.J.M.; Visualization, J.P.C. and J.A.M; Supervision, J.P.C., J.A.M, and D.J.M.; Funding Acquisition, J.P.C.and D.J.M.

## Acknowledgments

We would like to acknowledge Drs. Abhinava Mishra, Lauren Penfield, Wei Dai Guangxia Miao, and Allsion Gabbert for thoughtful discussions during the course of this work. We kindly thank Dr. David Bilder and Dr. Christopher Doe for *Drosophila* Scrib antibody and Dr. Jocelyn McDonald for Par-1 antibody. We acknowledge the Bloomington Drosophila Stock Center and Vienna Drosophila Resource Center for fly stocks. This work was funded by an American Cancer Society award PF-17-024-01-CSM to J.P.C and an National Institutes of Health award GM46425 to D.J.M.

---

**Supplementary figure 1:**
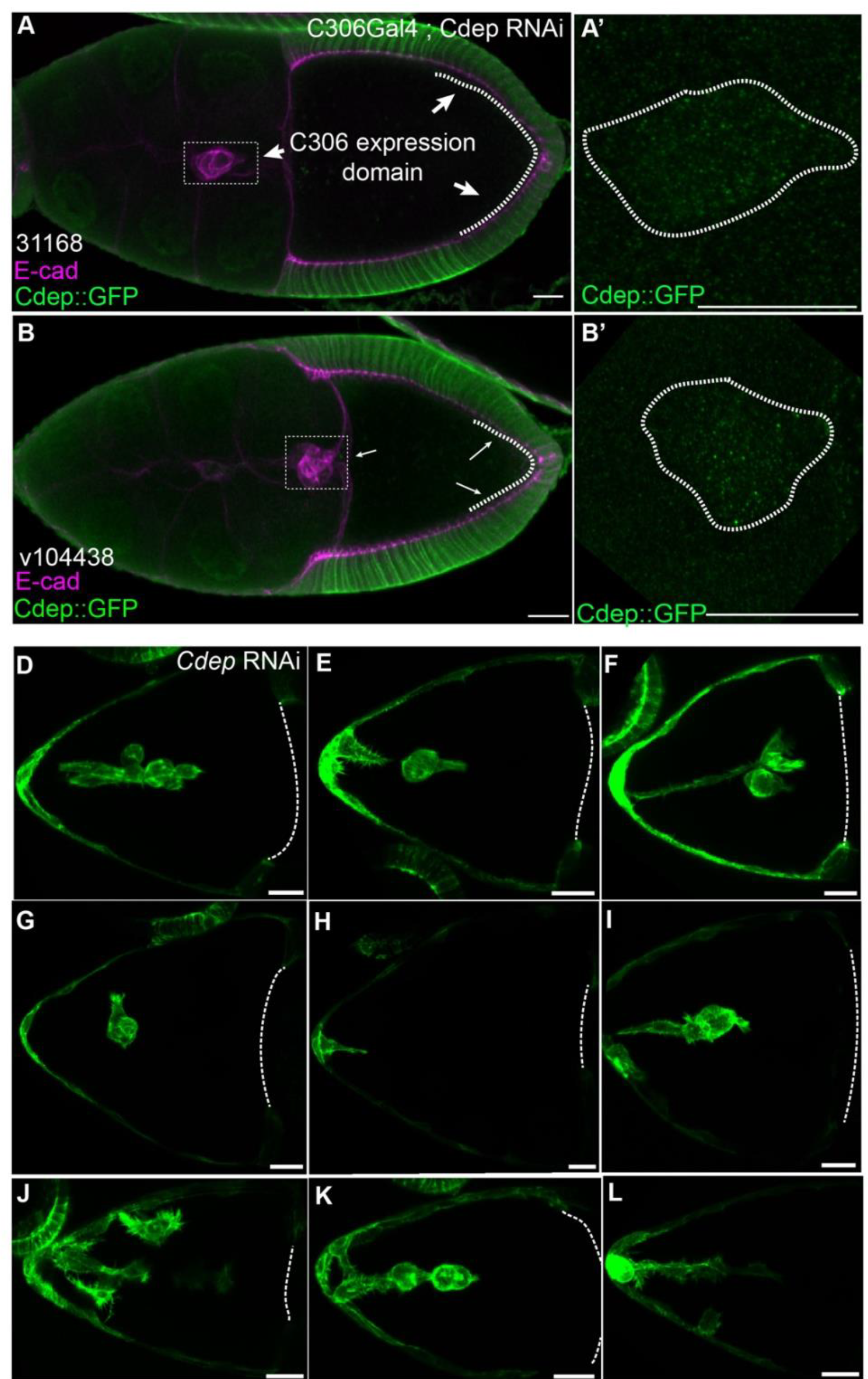
Cdep RNAi knocks down Cdep-GFP leading to cohesion defects. (A-B) Projections showing efficacy of Cdep knockdown in both the border cells (A,B - dashed box, A’ and B’) and in the epithelial cells that also are within the C306 expression domain (A,B - dashed line). (D-L) Representative images from individual time lapse movies showing morphological defects observed during live imaging after Cdep knockdown. All scale bars are 20 µm

**Supplemental Figure 2:**
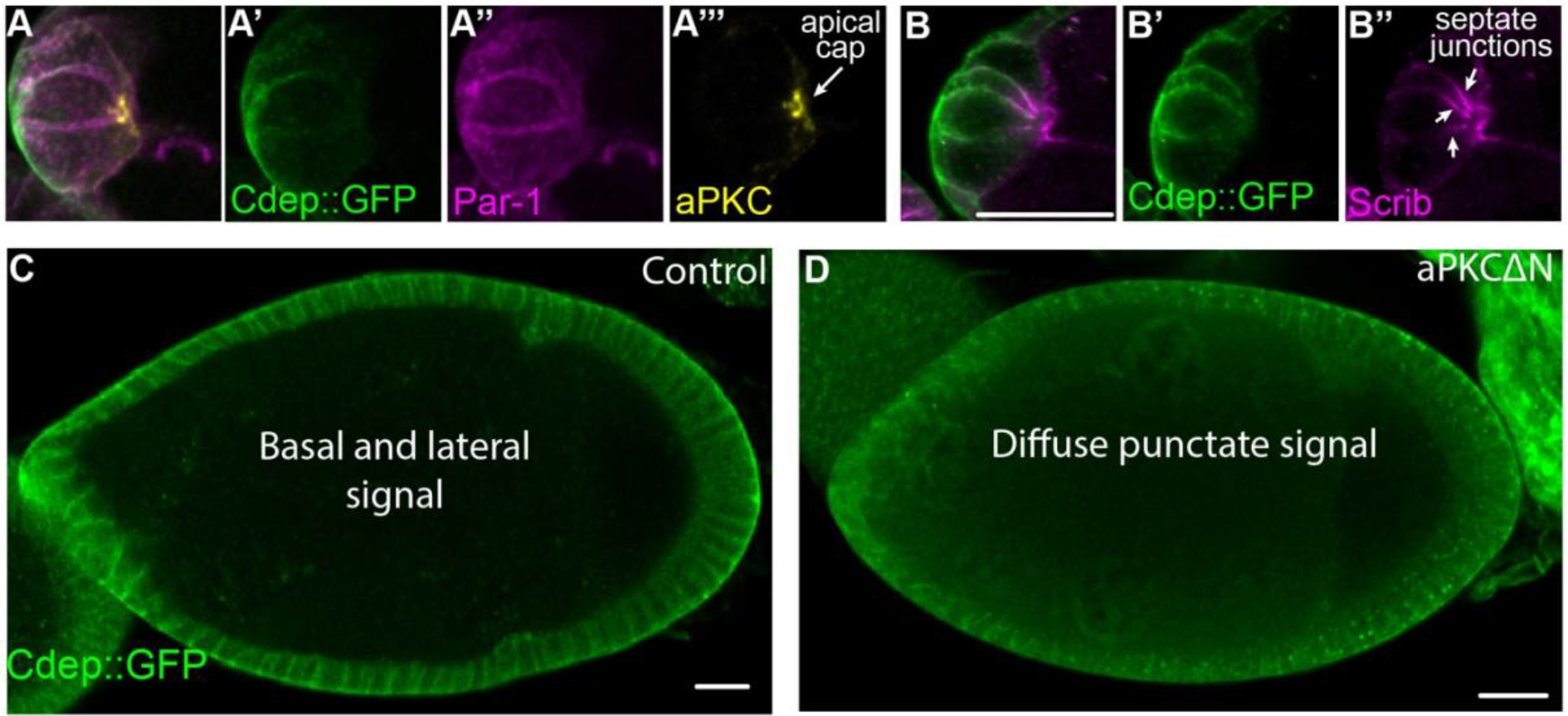
Cdep::GFP is asymmetrically localized to basolateral membranes. (A-A’’’) Composite image and individual channels from Fig 4H. (A’-A’’) Cdep::GFP (green) and Par-1 (magenta) antibody localizes to basal-lateral membrane while (A’’’) aPKC antibody (yellow) localizes to the apical cap of the polar cells. (B-B’’) Composite image and individual channels from Fig 4I. (B’) Cdep::GFP (green) localizes to basal-lateral membrane while (B’’) Scrib antibody localizes most strongly to sub-apical junctions at cell-cell contacts. (C) Cdep::GFP localizes to the basal-lateral membrane in all epithelial cells. (D) Upon expression of aPKC-delta-N Cdep loses its strong membrane localization, becoming more punctate and diffuse. All scale bars= 20 µm.

**Supplementary Figure 3.**
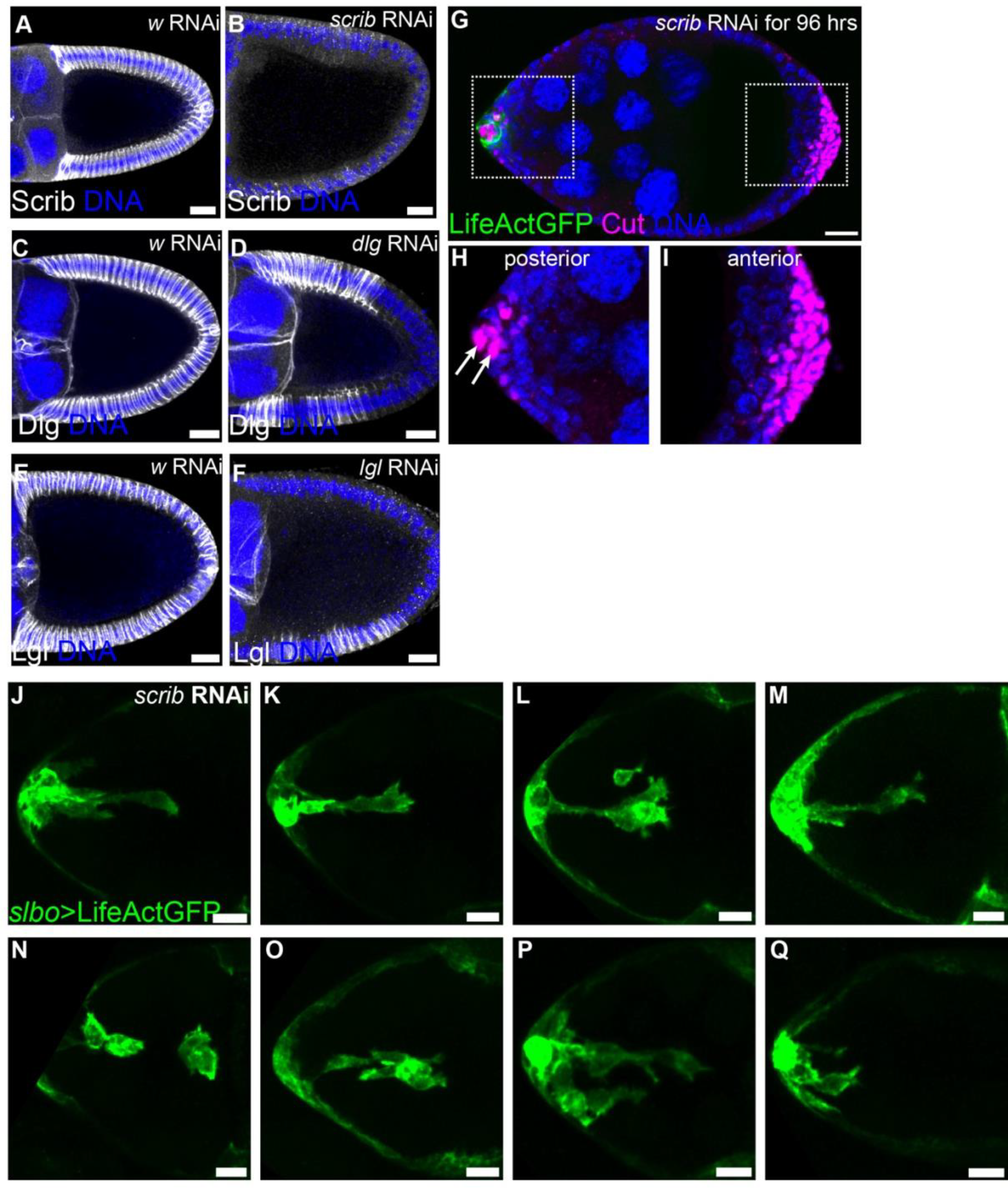
RNAi targetingScribble complex proteins reduces protein expression in egg chambers. (A-F) Scrib module protein knockdowns in posterior follicle cells (gray) after expressing *w, scrib, dlg*, or *lgl* RNAi. (G-I) Representative early stage 9 egg chamber expressing *scrib* RNAi for 96 hrs stained for *cut* (magenta) and DNA (blue). (H) Zoom of the anterior. Polar cell nuclei marked with arrows. (I) Zoom of the posterior. (J-Q) Representative images from individual time lapse movies showing morphological defects observed during live imaging of Scrib knockdown.

## References

Aceto, N., Bardia, A., Miyamoto, D.T., Donaldson, M.C., Wittner, B.S., Spencer, J.A., Yu, M., Pely, A., Engstrom, A., Zhu, H., et al. (2014). Circulating tumor cell clusters are oligoclonal precursors of breast cancer metastasis. Cell 158, 1110–1122.

Au, S.H., Storey, B.D., Moore, J.C., Tang, Q., Chen, Y.-L., Javaid, S., Sarioglu, A.F., Sullivan, R., Madden, M.W., O’Keefe, R., et al. (2016). Clusters of circulating tumor cells traverse capillary-sized vessels. Proc Natl Acad Sci USA 113, 4947–4952.

Betschinger, J., Mechtler, K., and Knoblich, J.A. (2003). The Par complex directs asymmetric cell division by phosphorylating the cytoskeletal protein Lgl. Nature 422, 326–330.

Bianco, A., Poukkula, M., Cliffe, A., Mathieu, J., Luque, C.M., Fulga, T.A., and Rørth, P. (2007). Two distinct modes of guidance signalling during collective migration of border cells. Nature 448, 362–365.

Bilder, D., Li, M., and Perrimon, N. (2000). Cooperative regulation of cell polarity and growth by Drosophila tumor suppressors. Science 289, 113–116.

Bonello, T.T., and Peifer, M. (2019). Scribble: A master scaffold in polarity, adhesion, synaptogenesis, and proliferation. J. Cell Biol. 218, 742–756.

Brumby, A.M., and Richardson, H.E. (2003). scribble mutants cooperate with oncogenic Ras or Notch to cause neoplastic overgrowth in Drosophila. EMBO J. 22, 5769–5779.

Cai, D., Chen, S.-C., Prasad, M., He, L., Wang, X., Choesmel-Cadamuro, V., Sawyer, J.K., Danuser, G., and Montell, D.J. (2014). Mechanical feedback through E-cadherin promotes direction sensing during collective cell migration. Cell 157, 1146–1159.

Cheung, K.J., and Ewald, A.J. (2016). A collective route to metastasis: Seeding by tumor cell clusters. Science 352, 167–169.

Cheung, K.J., Padmanaban, V., Silvestri, V., Schipper, K., Cohen, J.D., Fairchild, A.N., Gorin, M.A., Verdone, J.E., Pienta, K.J., Bader, J.S., et al. (2016). Polyclonal breast cancer metastases arise from collective dissemination of keratin 14-expressing tumor cell clusters. Proc Natl Acad Sci USA 113, E854–63.

Conod, A., Silvano, M., and Ruiz I Altaba, A. (2022). On the origin of metastases: Induction of pro-metastatic states after impending cell death via ER stress, reprogramming, and a cytokine storm. Cell Rep. 38, 110490.

Cooke, M., Kreider-Letterman, G., Baker, M.J., Zhang, S., Sullivan, N.T., Eruslanov, E., Abba, M.C., Goicoechea, S.M., García-Mata, R., and Kazanietz, M.G. (2021). FARP1, ARHGEF39, and TIAM2 are essential receptor tyrosine kinase effectors for Rac1-dependent cell motility in human lung adenocarcinoma. Cell Rep. 37, 109905.

Dai, W., Guo, X., Cao, Y.S., Mondo, J.A., Campanale, J.P., Montell, B.J.,Burrous, H., Streichan, S., Gov, N., Rappel, W.-J., et al. (2020). Tissue topography steers migrating Drosophila border cells. BioRxiv.

Duchek, P., Somogyi, K., Jékely, G., Beccari, S., and Rørth, P. (2001). Guidance of cell migration by the Drosophila PDGF/VEGF receptor. Cell 107, 17–26.

Fernández-Espartero, C.H., Ramel, D., Farago, M., Malartre, M., Luque, C.M., Limanovich, S., Katzav, S., Emery, G., and Martín-Bermudo, M.D. (2013). GTP exchange factor Vav regulates guided cell migration by coupling guidance receptor signalling to local Rac activation. J. Cell Sci. 126, 2285–2293.

Friedl, P., and Gilmour, D. (2009). Collective cell migration in morphogenesis, regeneration and cancer. Nat. Rev. Mol. Cell Biol. 10, 445–457.

Friedl, P., Sahai, E., Weiss, S., and Yamada, K.M. (2012). New dimensions in cell migration. Nat. Rev. Mol. Cell Biol. 13, 743–747.

Fulga, T.A., and Rørth, P. (2002). Invasive cell migration is initiated by guided growth of long cellular extensions. Nat. Cell Biol. 4, 715–719.

Geisbrecht, E.R., and Montell, D.J. (2004). A role for Drosophila IAP1-mediated caspase inhibition in Rac-dependent cell migration. Cell 118, 111–125.

Godt, D., and Tepass, U. (2009). Breaking a temporal barrier: signalling crosstalk regulates the initiation of border cell migration. Nat. Cell Biol. 11, 536–538.

Goode, S., and Perrimon, N. (1997). Inhibition of patterned cell shape change and cell invasion by Discs large during Drosophila oogenesis. Genes Dev. 11, 2532–2544.

Goode, S., Wei, J., and Kishore, S. (2005). Novel spatiotemporal patterns of epithelial tumor invasion in Drosophila discs large egg chambers. Dev. Dyn. 232, 855–864.

Haney, S., Konen, J., Marcus, A.I., and Bazhenov, M. (2018). The complex ecosystem in non small cell lung cancer invasion. PLoS Comput. Biol. 14, e1006131.

Hariharan, I.K., and Bilder, D. (2006). Regulation of imaginal disc growth by tumor-suppressor genes in Drosophila. Annu. Rev. Genet. 40, 335–361.

Harmansa, S., Alborelli, I., Bieli, D., Caussinus, E., and Affolter, M. (2017). A nanobody-based toolset to investigate the role of protein localization and dispersal in Drosophila. ELife 6.

Harrison, D.A., and Perrimon, N. (1993). Simple and efficient generation of marked clones in Drosophila. Curr. Biol. 3, 424–433.

Igaki, T., Pagliarini, R.A., and Xu, T. (2006). Loss of cell polarity drives tumor growth and invasion through JNK activation in Drosophila. Curr. Biol. 16, 1139–1146.

Kardash, E., Bandemer, J., and Raz, E. (2011). Imaging protein activity in live embryos using fluorescence resonance energy transfer biosensors. Nat. Protoc. 6, 1835–1846.

Khalil, A.A., and de Rooij, J. (2019). Cadherin mechanotransduction in leader-follower cell specification during collective migration. Exp. Cell Res. 376, 86–91.

Libanje, F., Raingeaud, J., Luan, R., Thomas, Z., Zajac, O., Veiga, J., Marisa, L., Adam, J., Boige, V., Malka, D., et al. (2019). ROCK2 inhibition triggers the collective invasion of colorectal adenocarcinomas. EMBO J. 38, e99299.

Lin, W.-H., Asmann, Y.W., and Anastasiadis, P.Z. (2015). Expression of polarity genes in human cancer. Cancer Inform. 14, 15–28.

Li, Q., Feng, S., Yu, L., Zhao, G., and Li, M. (2011). Requirements of Lgl in cell differentiation and motility during Drosophila ovarian follicular epithelium morphogenesis. Fly (Austin) 5, 81–87.

Lyda, J.K., Tan, Z.L., Rajah, A., Momi, A., Mackay, L., Brown, C.M., and Khadra, A. (2019). Rac activation is key to cell motility and directionality: An experimental and modelling investigation. Comput. Struct. Biotechnol. J. 17, 1436–1452.

Manseau, L., Baradaran, A., Brower, D., Budhu, A., Elefant, F., Phan, H., Philp, A.V., Yang, M., Glover, D., Kaiser, K., et al. (1997). GAL4 enhancer traps expressed in the embryo, larval brain, imaginal discs, and ovary of drosophila. Developmental Dynamics.

Mishra, A.K., Campanale, J.P., Mondo, J.A., and Montell, D.J. (2019a). Cell interactions in collective cell migration. Development 146.

Mishra, A.K., Mondo, J.A., Campanale, J.P., and Montell, D.J. (2019b). Coordination of protrusion dynamics within and between collectively migrating border cells by myosin II. Mol. Biol. Cell 30, 2490–2502.

Montell, D.J., Yoon, W.H., and Starz-Gaiano, M. (2012). Group choreography: mechanisms orchestrating the collective movement of border cells. Nat. Rev. Mol. Cell Biol. 13, 631–645.

Murphy, A.M., and Montell, D.J. (1996). Cell type-specific roles for Cdc42, Rac, and RhoL in Drosophila oogenesis. J. Cell Biol. 133, 617–630.

Ng, M.R., Besser, A., Danuser, G., and Brugge, J.S. (2012). Substrate stiffness regulates cadherin-dependent collective migration through myosin-II contractility. J. Cell Biol. 199, 545–563.

Nobes, C.D., and Hall, A. (1995). Rho, rac, and cdc42 GTPases regulate the assembly of multimolecular focal complexes associated with actin stress fibers, lamellipodia, and filopodia. Cell 81, 53–62.

Norden, C., and Lecaudey, V. (2019). Collective cell migration: general themes and new paradigms. Curr. Opin. Genet. Dev. 57, 54–60.

Pagliarini, R.A., and Xu, T. (2003). A genetic screen in Drosophila for metastatic behavior. Science 302, 1227–1231.

Parri, M., and Chiarugi, P. (2010). Rac and Rho GTPases in cancer cell motility control. Cell Commun. Signal. 8, 23.

Pinheiro, E.M., and Montell, D.J. (2004). Requirement for Par-6 and Bazooka in Drosophila border cell migration. Development 131, 5243–5251.

Qin, L., Yang, D., Yi, W., Cao, H., and Xiao, G. (2021). Roles of leader and follower cells in collective cell migration. Mol. Biol. Cell 32, 1267–1272.

Ramel, D., Wang, X., Laflamme, C., Montell, D.J., and Emery, G. (2013). Rab11 regulates cell-cell communication during collective cell movements. Nat. Cell Biol. 15, 317–324.

Revenu, C., Streichan, S., Donà, E., Lecaudey, V., Hufnagel, L., and Gilmour, D. (2014). Quantitative cell polarity imaging defines leader-to-follower transitions during collective migration and the key role of microtubule-dependent adherens junction formation. Development 141, 1282–1291.

Ridley, A.J. (2015). Rho GTPase signalling in cell migration. Curr. Opin. Cell Biol. 36, 103–112.

Rørth, P. (2009). Collective cell migration. Annu. Rev. Cell Dev. Biol. 25, 407–429.

Santoni, M.-J., Kashyap, R., Camoin, L., and Borg, J.-P. (2020). The Scribble family in cancer: twentieth anniversary. Oncogene 39, 7019–7033.

Scarpa, E., and Mayor, R. (2016). Collective cell migration in development. J. Cell Biol. 212, 143–155.

SenGupta, S., Parent, C.A., and Bear, J.E. (2021). The principles of directed cell migration. Nat. Rev. Mol. Cell Biol. 22, 529–547.

Stuelten, C., Parent, C.A., and Montell, D.J. (2017). Cell Motility in Cancer Metastasis: Mechanistic Insights from Simple Model Organisms. Nature Reviews Cancer.

Szafranski, P., and Goode, S. (2004). A Fasciclin 2 morphogenetic switch organizes epithelial cell cluster polarity and motility. Development 131, 2023–2036.

Theveneau, E., and Linker, C. (2017). Leaders in collective migration: are front cells really endowed with a particular set of skills? [version 1; peer review: 2 approved]. F1000Res. 6, 1899.

Wang, H., Qiu, Z., Xu, Z., Chen, S.J., Luo, J., Wang, X., and Chen, J. (2018). aPKC is a key polarity determinant in coordinating the function of three distinct cell polarities during collective migration. Development 145.

Wang, X., He, L., Wu, Y.I., Hahn, K.M., and Montell, D.J. (2010). Light-mediated activation reveals a key role for Rac in collective guidance of cell movement in vivo. Nat. Cell Biol. 12, 591–597.

Wu, M., Pastor-Pareja, J.C., and Xu, T. (2010). Interaction between Ras(V12) and scribbled clones induces tumour growth and invasion. Nature 463, 545–548.

Zegers, M.M., and Friedl, P. (2014). Rho GTPases in collective cell migration. Small GTPases 5, e28997.

Zhang, J., Goliwas, K.F., Wang, W., Taufalele, P.V., Bordeleau, F., and Reinhart-King, C.A. (2019). Energetic regulation of coordinated leader-follower dynamics during collective invasion of breast cancer cells. Proc Natl Acad Sci USA 116, 7867–7872.

Zhao, M., Szafranski, P., Hall, C.A., and Goode, S. (2008). Basolateral junctions utilize warts signaling to control epithelial-mesenchymal transition and proliferation crucial for migration and invasion of Drosophila ovarian epithelial cells. Genetics 178, 1947–1971.

